# Novel modular pressure and flow rate balanced microfluidic serial dilution networks on printed circuit boards: Designs, Simulations and Fabrication

**DOI:** 10.1101/270124

**Authors:** Nikolaos Vasilakis, Konstantinos I. Papadimitriou, Hywel Morgan, Themistoklis Prodromakis

## Abstract

Fast, efficient and more importantly accurate serial dilution is a requirement for many chemical and biological microfluidic-based applications. Over the last decade, a large number of microfluidic devices has been proposed, each demonstrating either a different type of dilution technique or complex system architectures based on various flow source combinations. In this work, a novel serial dilution architecture is demonstrated, implemented on a commercially fabricated printed circuit board (PCB). The proposed single layer, stepwise serial diluter comprises an optimised microfluidic network, where identical dilution ratio per stage can be ensured, either by applying equal pressure or equal flow rates at both inlets. The advantages of the proposed serial diluter are twofold. Firstly, it is structured as a modular unit cell, simplifying the required fluid driving mechanism to a single source for both sample and buffer solution. Thus, this unit cell can be seen as a fundamental microfluidic building block, which can form multistage serial dilution cascades, once combined appropriately with itself or other similar unit cells. Secondly, the serial diluter has been fabricated entirely using commercial PCB technologies, allowing the device to be interfaced with standard electronic components, if more complex miniature point-of-care (PoC) systems are desired, where the small footprint and accuracy of the device is of paramount importance.

## 1 Introduction

Almost every quantitative chemical and biological assay relies heavily upon accurate and prompt serial dilution. Its usefulness lies in the fact that with a suitable sample preparation procedure, a wide variety of known scaled concentration samples can be generated using a suitable diluent [1]. For example, widely known quantitative assays, such as real time polymerase chain reaction (q-PCR) or antigen assays, e.g. Enzyme-Linked Immunosorbent Assay (ELISA), require serial dilutions of a known analyte to produce several reference calibration signals which can be later used for standard or calibration curves. Over the last 15 years, many continuous flow microfluidic dilution devices have been proposed for the generation of multiple stepwise concentrations or concentration gradients with logarithmic [2, 3] or linear dilution [4–7]. However, the majority often requires multiple flow sources (pressure or flow rate) to be employed for driving fluids through the device. Their operation could be potentially unstable and they lack ease of use [8] and integration with other components in a monolithic manner. To tackle the above, herein, a single layer, stepwise serial diluter is proposed comprising an optimised microfluidic network, where identical dilution ratios can be ensured either by applying equal pressure or equal flow rates at both inlets. It is a modular unit-cell that simplifies the fluid driving mechanism to a single source for both sample and buffer. Finally, this unit cell can be seen as a fundamental microfluidic building block, forming multistage serial dilution cascades, once combined with itself or other similar unit cells.

Microfluidic devices are particularly suited for PoC diagnostics, and for these devices one route to development is to adopt standard assays from the laboratory or “assay track” to create diagnostic kits. There are now several PoC platforms that perform *sample-to-answer* operations [9]. Although they are able to deliver multiple functions in a monolithic manner, meeting the “ASSURED” criteria [10] (**A**ffordable, **S**ensitive, **S**pecific, **U**ser friendly, **R**obust and **R**apid, **E**quipment-free, and **D**eliverable to end-users) proposed by WHO is always a challenge [9, 11]. This implies that a successful microfluidic PoC device needs, besides sample and reagents manipulation, to be able to perform the detection and signal processing on chip in the most cost- and space-effective manner possible. An alternative microfluidic system based entirely on PCB manufacturing technology is demonstrated in this work: The lab on printed circuit board (LoPCB). As a concept was initially proposed by Lammerink *et al.* [12] and named mixed circuit board (MCB). This technology was further developed by Merkel *et al.* [13], who integrated microfluidic platforms combined with electronic components by adding processing steps in the PCB fabrication technology. In detail, their manufactured prototypes comprised standard PCB features such as copper tracks, combined with electronic assemblies, microfluidics and sensing electrodes, all fully integrated in a monolithic manner towards a complete, functional LoPCB device. Using the same approach various devices have been demonstrated, such as a pH regulation system comprising an optical sensor and a CO_2_ diffuser [14] as well as a 4-layer PCB self-filling micropump [15]. Moreover, a valve-less micropump comprising piezoelectric discs and diffuser/nozzle components has been also reported [16]. More recently micromixers [17], nucleic acid amplification chips [18] and chemical sensors [19] have been demonstrated. Electrochemical biosensing on PCB requires gold (Au) or silver (Ag) plated sensing pads integrated with microfluidic networks for reagent manipulation and sample preparation on the same platform [20, 21]. The interested reader can find a thorough demonstration of the potentials of LoPCB technology here [22].

From all the above, it becomes evident that a major advantage of the LoPCB technology is that it provides an analytical platform, where the microfluidic components for sample preparation and reagent manipulation can be integrated with electrochemical sensors and bespoke circuitry. Since there is no need for separate electronics and assay platforms, the required footprint of the whole measuring setup decreases, providing direct, more efficient electrochemical sensing in smaller areas/volume, contrary to, for example, bulky high-sensitivity spectrometric apparatus [23]. Furthermore, the combination of both, biochemistry and appropriate electronics on the same platform reduces noise interference from various interconnection solutions and as a result improves the measurement’s signal-to-noise ratio (SNR) [24]. Finally, the high degree of electronics integration, the exceptional precision, and the accumulated experience and skills of a mature industrial manufacturing process highlight the great advantage to LoPCB platforms [25, 26]. A detailed architecture of a standard “3-layer” LoPCB device, with integrated microfluidics, biosensors and electronics is shown in Figure 1. The inner bottom layer is a photo-lithographically patterned dry photoresist laminated on an FR-4 substrate which can be modified, depending on the application, and forms the integrated microfluidic network. The key aspect of our technology is the integration of microfluidic components with electronics, and biosensing electrodes to create a monolithic device as illustrated conceptually in Figure 1. Taking into consideration that a PCB based PoC testing platform implementing a quantitative bioassay requires a microfluidic network that generates samples of known concentrations to acquire a standard curve on chip, i.e. a microfluidic serial diluter, it is evident that an integrated microfluidic diluter on PCB should minimise the required flow sources number while tolerating flow source instabilities resulting in a robust operation and affordable implementation.

There are two main categories of diluters, the diffusion-based and the multi-step diluters. The diffusion-based designs [27, 28] exploit the concentration gradient formed along a microfluidic channel to create samples of different concentrations across the same microchannel. The main challenge for these devices is to obtain linear concentration gradients, due to the inherent non-linear diffusion mechanism [29]. Multi-step microfluidic dilution devices were firstly proposed by Jacobson *et al* in 1999 but the resulting dilution ratio was neither linear nor logarithmic [30].

In 2001 Whitesides’ group proposed the first linear concentration gradient device [31], while two years later an improved device following the same approach, including staggered herringbone micromixers, was demonstrated [32]. Later, Kim *et al.* developed a serial dilution microfluidic chip that produced logarithmic and linear step-wise concentrations, respectively. However, the inlet flow rates of both buffer and sample were not equal. The logarithmic device required a buffer-to-sample ratio of 1:36 (sample 0.5 mL/h vs buffer 18.0 mL/h) while the linear one needed a 1:2.75 ratio (sample 1.0 mL/h vs buffer 2.75 mL/h) [29]. Lee *et al.* developed a microfluidic network-based device consisting of three different layers. This design could combine three different samples with a buffer solution. The buffer solution diluted the samples in a ratio 1:4 and through the 3-layer microfluidic network all possible combinations were formed in the output device ports. Nonetheless, the buffer flow rate was four times the flow rate of each sample [33]. Recently, this principle was enhanced by adjusting not only the channel length but also the channel width. In 2013, Weibull *et al.* reported a stepwise dilution generator, including logarithmic and linear gradients in a two-layer microfluidic implementation. Although two different dilutions gradients were implemented on the same device, they have separated inlets, microfluidic network and outlets. Each device was able to generate only three dilution ratios [34]. More recently, Occhetta *et al.* developed a high-throughput microfluidic screening platform generating six different concentration outflows aimed at 3D cell cultures. Two different variations of linear and logarithmic dilution were created using network channels length as the only variable. The achieved flow rate was 24 μL/h (12 μL/h through every inlet) through the linear dilution device and 45 μL/m through the logarithmic. It is notable that the logarithmic dilution device required a flow ratio between sample and buffer inlets of 1:3.5 [35]. The majority of the aforementioned microfluidic implementations required multiple sources (syringe pumps) and consequently led to complicated final system architectures.

**Figure 1:**
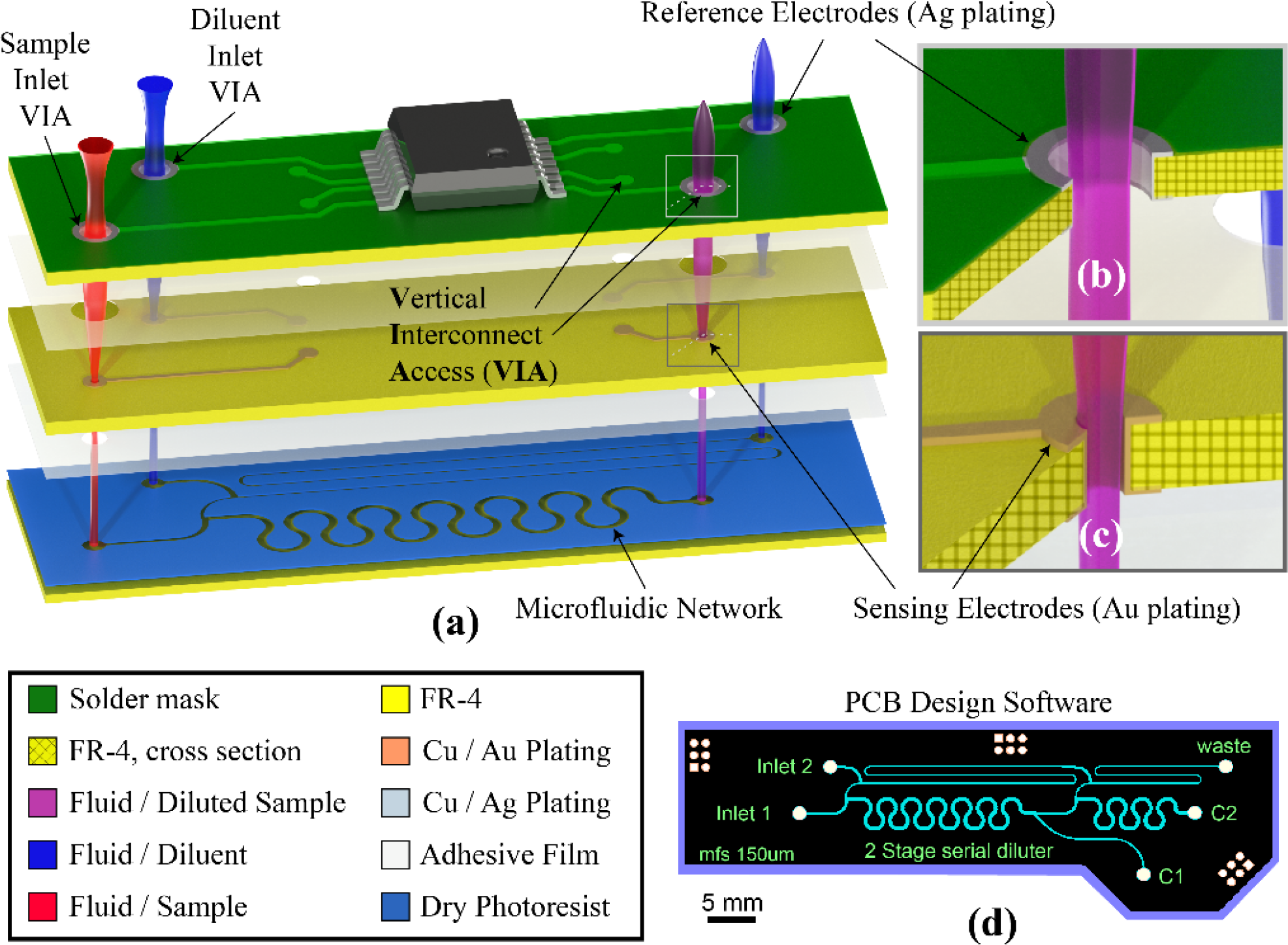
The LoPCB. a) Exploded view of the integrated design b) and c) cross sectional views of the VIAs employed as reference and sensing electrodes, respectively d) the PCB CAD design of the microfluidic layer used for the fabrication of the samples using commercially available technology.

Herein, we describe a simple, yet elegant, single layer, stepwise serial diluter that could address the requirements of an integrated LoPCB system for quantitative PoC applications. The flow rate through the diluter allows short assay duration time, i.e. less than 5 minutes. The unit cell is optimised to generate identical dilution rate when, a) either the applied pressure at the inlets is the same or b) the flow rates through both inlets are equal. Therefore, the architecture revolves around pressure and flow rate balanced design. It can also be considered as a modular unit cell that simplifies the fluid input driver to a single source for both, sample and diluent solutions. The dilution ratio remains stable, with tolerance to the input pump instabilities. Hence, a single syringe pump or even a compressed air chamber (e.g. blister packaging) could be employed, reducing the complexity, cost and overall weight and footprint of the final device. This novel pressure and flowrate balanced device could be the ideal candidate for quantitative PoC tests and may be used as a building block for multistage serial dilution cascade sample preparation components generating diluted samples of known concentrations to derive calibration curves on chip.

## 2. Materials and methods

### 2.1 Pressure and flow rate balanced unit cell design

Figure 2 illustrates the core design principle of the pressure and flow rate balanced unit cell. The two inlets A and B (see Figure 2a) are designed to be co-linear in pairs with the two outlets (C and D) to make potential ladder designs easier to be aligned during laid-out phase. Inlets A and B serve either as sample or diluent inlets according to the desired dilution ratio (in this case *C*_*c*_ = 2: 3 *C*_*B*_ or 1: 3 *C*_*A*_, respectively). As shown in Figure 2a, the design is divided into four zones. Zones 1 (between points 4 and 3) and 2 (between points 1 and 2) are the device inlets. Zone 3 starts after the branching point 2, where the inlet A stream splits into two sub-streams, and ends at point 6 (outlet D). The hydraulic resistance of zone 3 is represented by the notation R_2−6_ in the electrical analogue of the design (see Figure 2b). Zone 4 starts at merging point 3, where sample and diluent streams merge and ends at point 5 (outlet C). The latter zone is a planar micro-mixing channel consisting of 6 double circular loops. Uniform mixing of the two reagents is imperative for the diluter exemplary performance.

**Figure 2:**
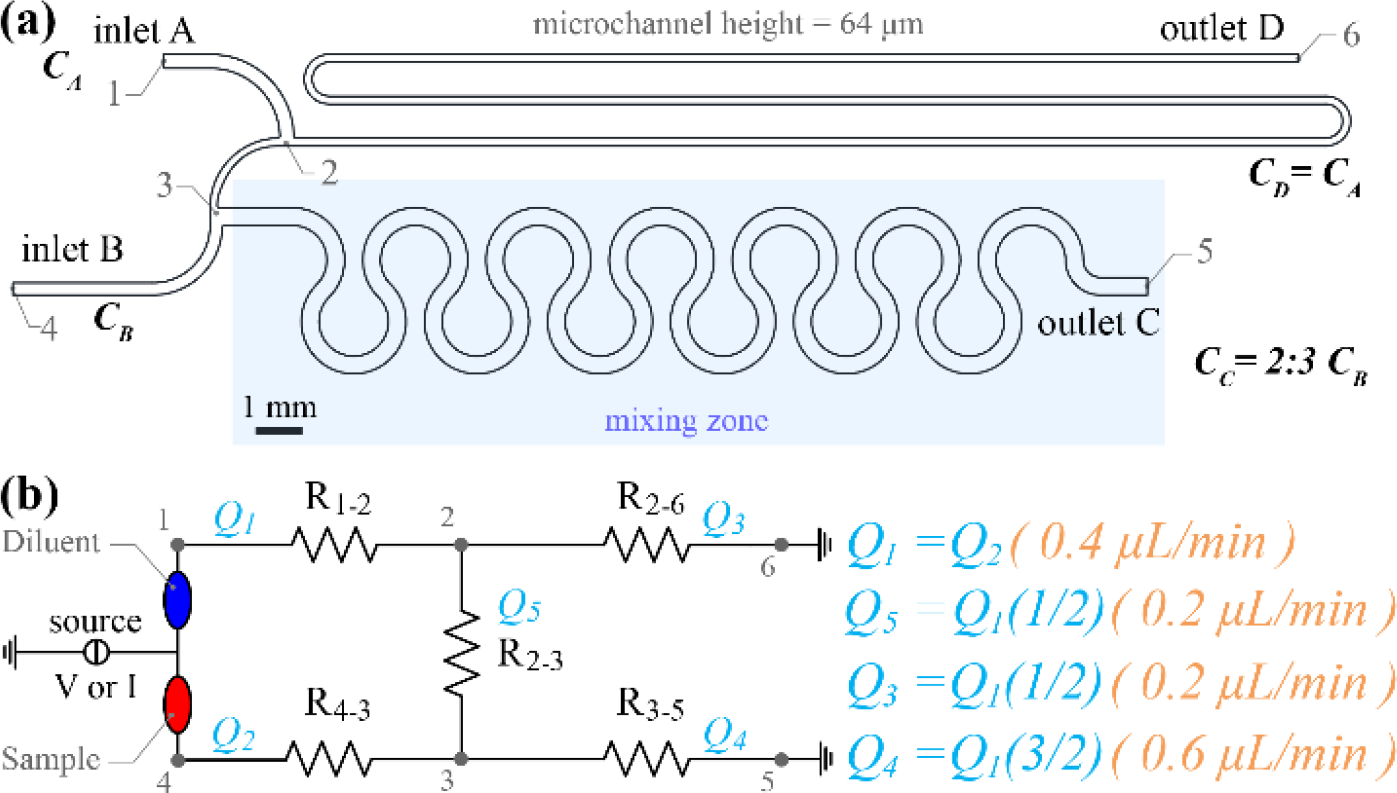
Serial diluter pressure and flow rate balanced unit cell. a) Optimised design generating 2:3 diluted sample b) microfluidic network hydraulic resistance electrical analogous.

Equation 1 can be used for a rough estimation of the required mixing microchannel length considering both, the flow rate and the diffusion coefficient constraints of the given application.

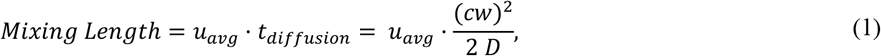

where *Mixing Length* is the estimated required length in metres, *u*_*avg*_ defines the average fluid velocity along the microchannel in m/s, *t*_*diffusion*_ is the required time for a molecule in the sample stream to diffuse across the channel, *cw* is the width of the microfluidic channel in metres, and *D* is the diffusion coefficient of the molecules in the buffer solution in m^2^/s. Equation 1 can be applied to the proposed design, since mixing is mainly driven by molecular diffusion because the Reynolds number is low (<100) and consequently the flow field is laminar. Furthermore, any additional convection driven mixing mechanism induced by the microchannels geometry will only lead to an enhancement of the mixing efficiency. Hence, the result of equation 1 can be always considered as an “*overestimation*” of the required mixing length.

Figure 2b presents the electrical analogue of the serial diluter unit cell design in Figure 2a. *Q*_1_ is the current (flow rate analogous) through R_1-2_ (inlet A) and *Q*_2_ the current through R_4-3_ (inlet B). As an initial step, the two flow rates (or currents) were assumed equal, in order to facilitate the dilution network operation using only one source. The hydraulic resistances R_1-2_, R_2-3_ etc. in this sketch are named after the numbered points illustrated in Figure 2a. The dilution ratio (DR) is determined by the flow rate ratio of the two streams joining at the merging point 3. An additional requirement for the design optimisation phase was that the pressure at the two inlets should be equal as well. This resulted in a pressure-balanced unit cell design exhibiting several advantages over the unbalanced ones described in the literature. More specifically, the novelty of the proposed design lies in the fact that any instabilities in the mixing ratio induced by input flow fluctuations (due to interfacing tubing, syringe pump vibrations etc.) will be mitigated, due to similar behaviour of device inlets with respect to pressure.

This pressure-balanced design will also be able to operate with a more cost-effective, low-power pneumatic pressure source (either pneumatic pump or compressed air chambers), instead of the traditional syringe pump to drive sample and diluent through the serial diluter. For example, in the case of a PoC handheld device, there would be no need for an accurate and precise syringe pump. In addition, the two inlets can be “bridged” (as illustrated in Figure 2b) utilising the same pressure source. In this case, any pressure variations at the inlets will not affect the mixing ratio in every unit cell but will only affect the total flow rate. This implies that the required interfacing ports for the sample and buffer manipulation can be reduced from 2 to only 1, reducing the overall size of the final device even more.

A serial dilution ladder network comprising two stages stemmed from the optimised design already shown in Figure 2a and can now be seen in Figure 3a, demonstrating the resulting 2-stage serial diluter design after simulation optimisation. Inlet flow rates A and B were again assumed to be equal. The DR for both stages has been selected to be 2:3, a reasonable value for many applications. The flow rate at both inlets of the second stage (point 6 and sampling point 5) is half compared to the flow rate through inlets A and B (see Figure 3b). Also, the length of the mixing microchannel between points 3’ and 5’ is shorter (approximately half) than the micro-mixing zone of Stage 1. The highlighted 1^st^ stage in Figure 3a and b is the balanced modular unit cell that can be used as building block for cascade designs forming *n-stage* serial diluters. However, this device feature is only valid for liquids (sample and diluent) of similar viscosity. In different case, the diluted sample flowing though branch point 5 will be of different viscosity and will affect the pressure drop balance between the microchannels of the 2^nd^ stage and consequently the resulting dilution ratio. It is also worth mentioning that every subsequent stage after the 1^st^ operates with half flow rate compared to the other ones, due to the design architecture and subsequently, the mixing efficiency of the micro-mixing zone is expected to be higher than the 1^st^ stage.

**Figure 3:**
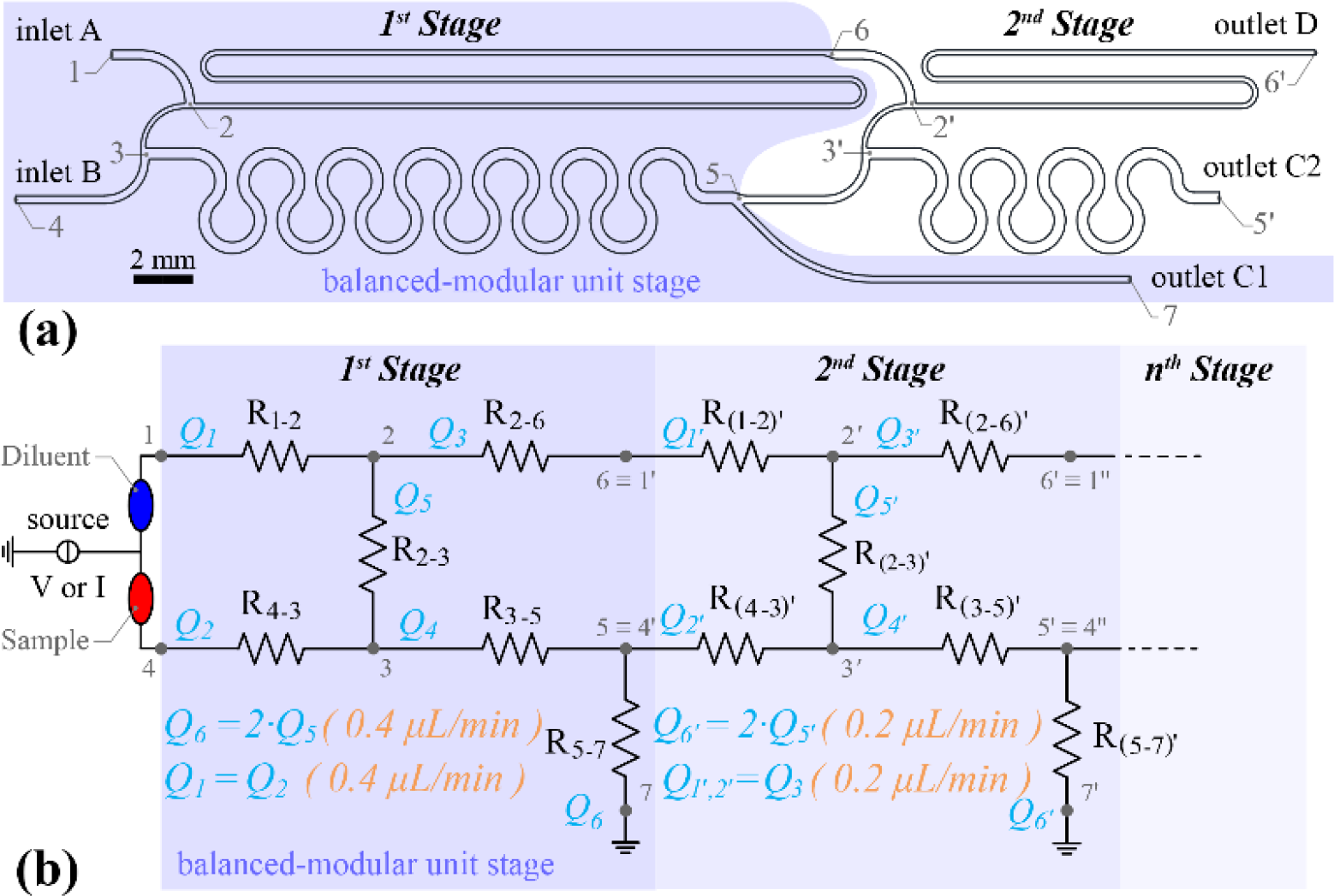
The 2-stage stepwise serial diluter a) design overview highlighting the balanced modular unit stage b) electrical network analogous of an N-stage serial diluter design.

### 2.2 Analytical design, simulation model and optimisation

It is quite common to design complex microfluidic networks based on simplified equations for hydraulic resistance. Such an approach is accurate only when the microchannel cross section is the same throughout the whole device. Microfluidic chips that are based on unique microchannel cross sections usually have very long channels, with high pressure drops and can suffer from flow instabilities, due to material deformation and capacitive effects. In this work, we optimised not only all the microchannel lengths but also the widths, based on the targeted flow rates. In our case, the use of simplified hydraulic resistance equations for rectangular microchannel is not recommended, due to the error from the actual value, which is typically around 20% [36]. A more accurate approximation based on an electric circuit analogy was used as an initial step for the designs, following the methodology described by [36]. The relationship between the volumetric flowrate *Q* and the pressure drop *ΔP* along a perfectly rectangular channel for a steady-state, pressure-driven, fully developed, laminar flow of an incompressible, uniform-viscous Newtonian liquid (aqueous solutions) is well known, assuming that the pressure gradient along the microchannel is uniform can be described by the following simplified Hagen-Poiseuille’s law:

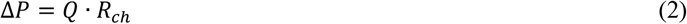

where *R*_*ch*_ is the hydraulic resistance of the microfluidic channel in Pa m^3^ s^−1^.

For a rectangular microchannel the hydraulic resistance can be calculated as the summation of a Fourier series [37], where the first six terms of the series are sufficient to calculate *R*_*ch*_, generating a negligible error of ~10^−6^:

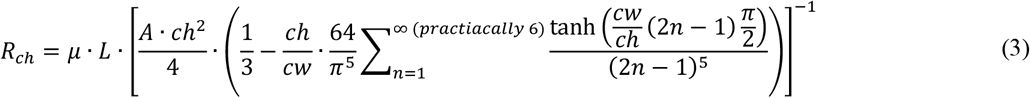

where *μ* is the dynamic viscosity of the liquid in s Pa ( *μ*_*water*_ = 10^−3^ *s · Pa*), *L* the length of the microfluidic channel, *ch* the channel height, *cw* the channel width and *A* the cross-sectional area of the microchannel (i.e. *A = ch · cw*).

Equation (2) can be viewed as a hydraulic analogous equation of Ohm’s law. Pressure drop *ΔP* is analogous to the voltage drop *ΔV*, flow rate to the current through the resistor and hydraulic resistance to the electrical resistance. A mass conservation equation in each node can be used, analogous to Kirchhoff’s current law (KCL). The required number of equations to define the hydraulic resistances of the microfluidic network can be defined using the energy conservation equations, again in perfect analogy with Kirchhoff’s voltage law (KVL). On the other hand, equation 3 indicates that the hydraulic resistance is a function of both channel width and length (channel height is defined by the thickness of the dry photoresist (DPR)). The final shape of the microfluidic device was defined after COMSOL Multiphysics^®^ optimisation simulations.

More specifically, using the resulting hydraulic resistances we defined channel widths and lengths based on the technology limitations (minimum feature size, MFS). The desired pressure drop along the unit cell to mitigate the interfacing challenges was an additional constraint. The initial topology of the device was constrained by geometrical factors to facilitate the cascade design and minimise the footprint of the device, as much as possible. The initial geometry was designed using a parametric 3D CAD software [38] while COMSOL Multiphysics^®^ [39] was used for the 3D modelling, simulation and optimisation of the proposed designs. The numerical model included laminar flow and transport of diluted species models and was solved in a sequential, coupled manner, as shown in previous work [40]. An automated iterative simulation methodology was employed where the parametric 3D model was transferred to the simulation software, that after deriving a converged solution of the flow and pressure field, amended the geometry parameters of the model feeding back the results to the 3D CAD software. This loop was repeated several times until the following two constrains were fulfilled:

1. the need to maintain the flow rates presented in Figures 2b and 3b at the inlets and outlets of the system
2. the constraint that the absolute average pressure difference between inlets A and B should be less than 0.01 Pa.

The diffusion coefficient of the sample in the buffer solution was assumed to be 6.67 10^−10^m^2^/s (Glucose in aqueous solutions) [41]. Both optimised designs (the balanced unit cell shown in Figure 2 and the 2-stage serial diluter in Figure 3) were transferred to Altium Designer [42], a commercially available software used for PCB design. The produced standard Gerber files were then submitted to an industrial PCB manufacturer, Newbury Electronics Ltd. for device fabrication.

#### Laminar flow model

A laminar incompressible steady state flow model has been used, since the studied volumetric flow rates along the microchannel was between 0.2 and 0.8 μL/min. The medium flowing through the channel via both inlets was selected to be water at 20°C. Considering the kinematic viscosity of the fluid, such flow rates have Reynolds number within the laminar range (Re<<1000). The model takes into consideration mass and momentum conservation equations for steady state incompressible flow, where gravitational effects are neglected. The equations can be seen below:

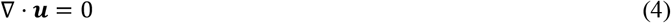

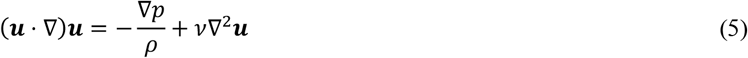

with ***u*** is the velocity vector, *p* the pressure, *ρ* the density of the medium and *v* the kinematic viscosity [43]. A parabolic velocity profile of fully developed laminar flow was formed over the two inlets. No-slip velocity on the channel walls was used as a boundary condition. The dilution networks were numerically investigated for the following flow rates: 0.2 μL/min, 0.4 μL/min, 0.6 μL/min and 0.8 μL/min.

#### Diffusion model

The concentration-based advection-diffusion model of COMSOL Multiphysics^®^ was employed for mixing efficiency studies. The mass balance equation describing the steady-state problem is described by the following relation:

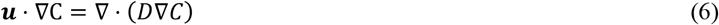

where *C* is the concentration of the diluted species in mol/L. The concentration of diluted species over inlets A and B, in both designs, was set equal to 0.00 and 1.00 mol/L, respectively, i.e. *C*_*A*_= 0.00 mol/L and *C*_*B*_ = 1.00 mol/L. At this point, the interested reader should note that any correlation between the fluid kinematic viscosity or density and the glucose concentration was neglected.

#### Mixing efficiency

Throughout the literature several micromixer performance quantification methods are proposed [44–46]. Since adequate mixing is defined as the homogeneity of the mixed components, the distribution of the concentration over the computational nodes of a flow cross-section can be used to evaluate the degree of mixing. Ngyen proposed the Mixing Efficiency (ME) [46] as a mixing quantification parameter with the following mathematical expression:

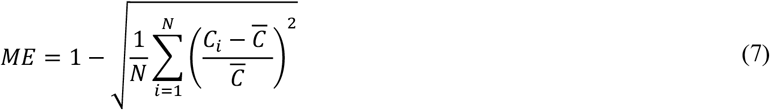

where 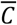 is the concentration of fully mixed medium (in our case 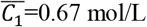 mol/L and 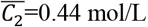 mol/L), *C*_*i*_ is the concentration at a given position (spatial discretisation point) and N is the total number of discretisation points over the outflow surface. In other words, equation 5 is a normalised standard deviation of the diluted species concentration over the computational nodes, with values ranging from 0.00 to 1.00 (0.00 defining completely unmixed species and 1.00 an ideally uniform mixed flow cross-section) [46]. In this work, the ME has been calculated on the computational nodes of the outflow surface.

### 2.3 Device fabrication

The commercially fabricated devices comprise a standard FR-4 sheet and a 64 μm thick DPR, patterned using the standard lithography process employed in commercial PCB manufacturing technology. Figure 4 illustrates the in house postprocessing of the final device which comprises the PCB based microfluidic layer on the bottom (FR-4 and DPR) and two additional layers of polymethyl methacrylate (PMMA) to create the microfluidics interface. The devices were sealed using polyethylene terephthalate (PET) sealing film (50 μm thickness, PET, VWR ^®^ Polyester Sealing Films for ELISA). After opening sample inlet and outlet holes on the PET film using a CO_2_ laser cutter (Epilog Mini 24 Legend Laser System, USA), both the PCB microfluidic network and the PET film were placed in a pouch laminator (60 C, lamination speed around 4 mm/s). Then the first PMMA interface layer (Figure 4, DS1 and PMMA 1) was created. A double sided adhesive 130 μm thick film (3M™ High Performance Acrylic Adhesive 200MP) was laminated on a 3 mm thick PMMA sheet (room temperature, lamination speed around 4 mm/s). Inlet and outlet ports were opened using again the laser cutter. Subsequently, both the sealed microfluidic device and the first interface layer were aligned and 40 kPa pressure applied for 1h to bond the two components together. The second interface layer was fabricated similarly. After opening the inlet ports and the outlet wells (laser cutter machining) the inlet ports were threaded for interfacing with commercial connectors. A 2-stage PMMA serial diluter designed and prototyped following the same principle but with much bigger footprint generating dilutions of 1:6 ratio is described in [47].

**Figure 4:**
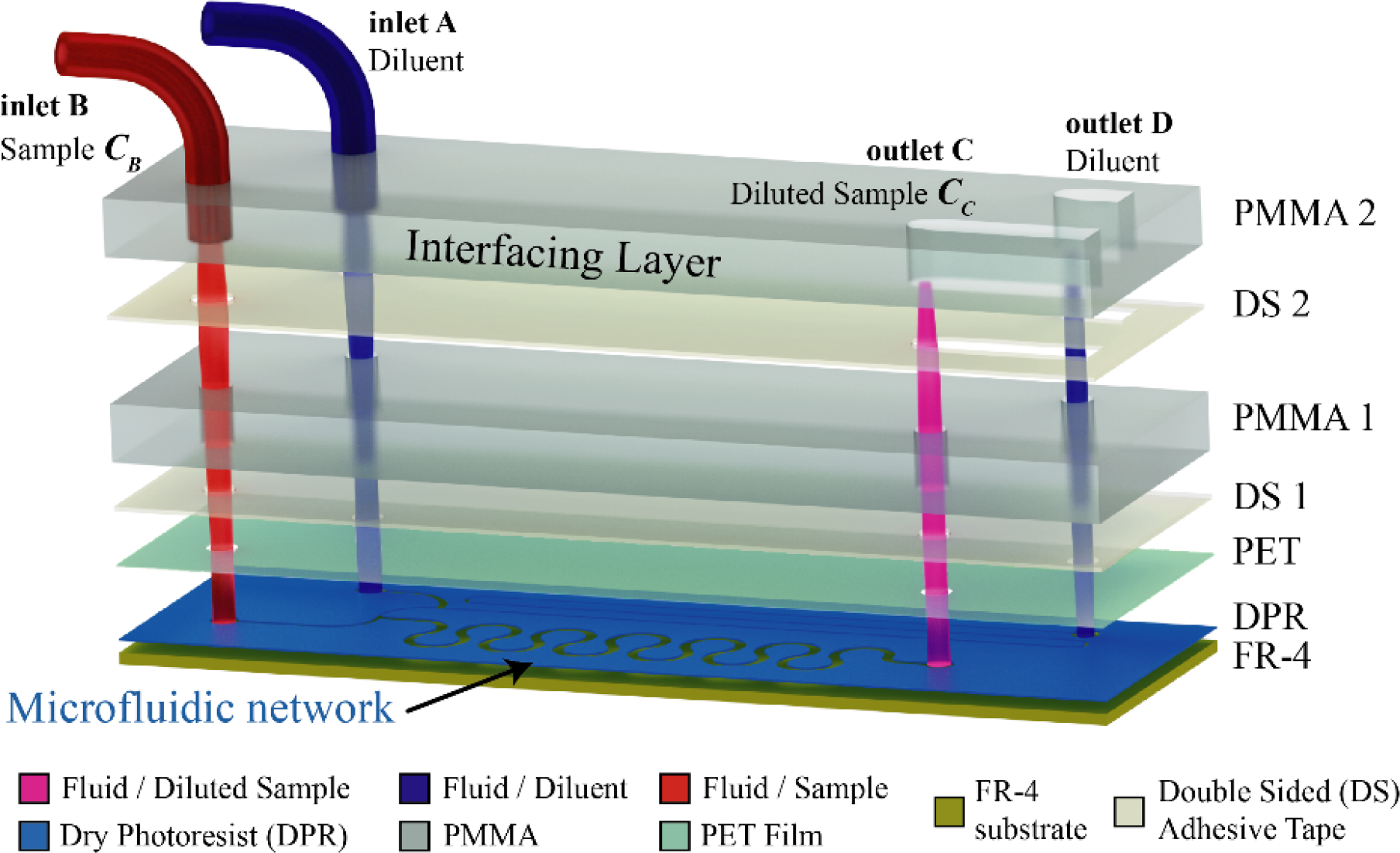
Exploded view presenting the various PMMA and PET layers used to interface the PCB fabricated prototype with the required tubing for the dilution performance characterisation experiments. Both outlets C and D supply the two wells with diluted sample and bypassing diluent respectively.

Figure 5a shows the experimental setup with one syringe pump to drive both diluent and sample through the 2-stage PCB serial diluter, while Figure 5b shows the top view of the prototype device. More specifically, Figure 5b shows that the three outlets of the 2-stage PCB diluter (i.e. C_1_, C_2_ and D) are driving the samples filling three wells formed in the PMMA sheet. The area of the three wells was designed to have the same ratio as the flow rate ratio through the three outlets to maintain equal pressure on the device outlets.

**Figure 5:**
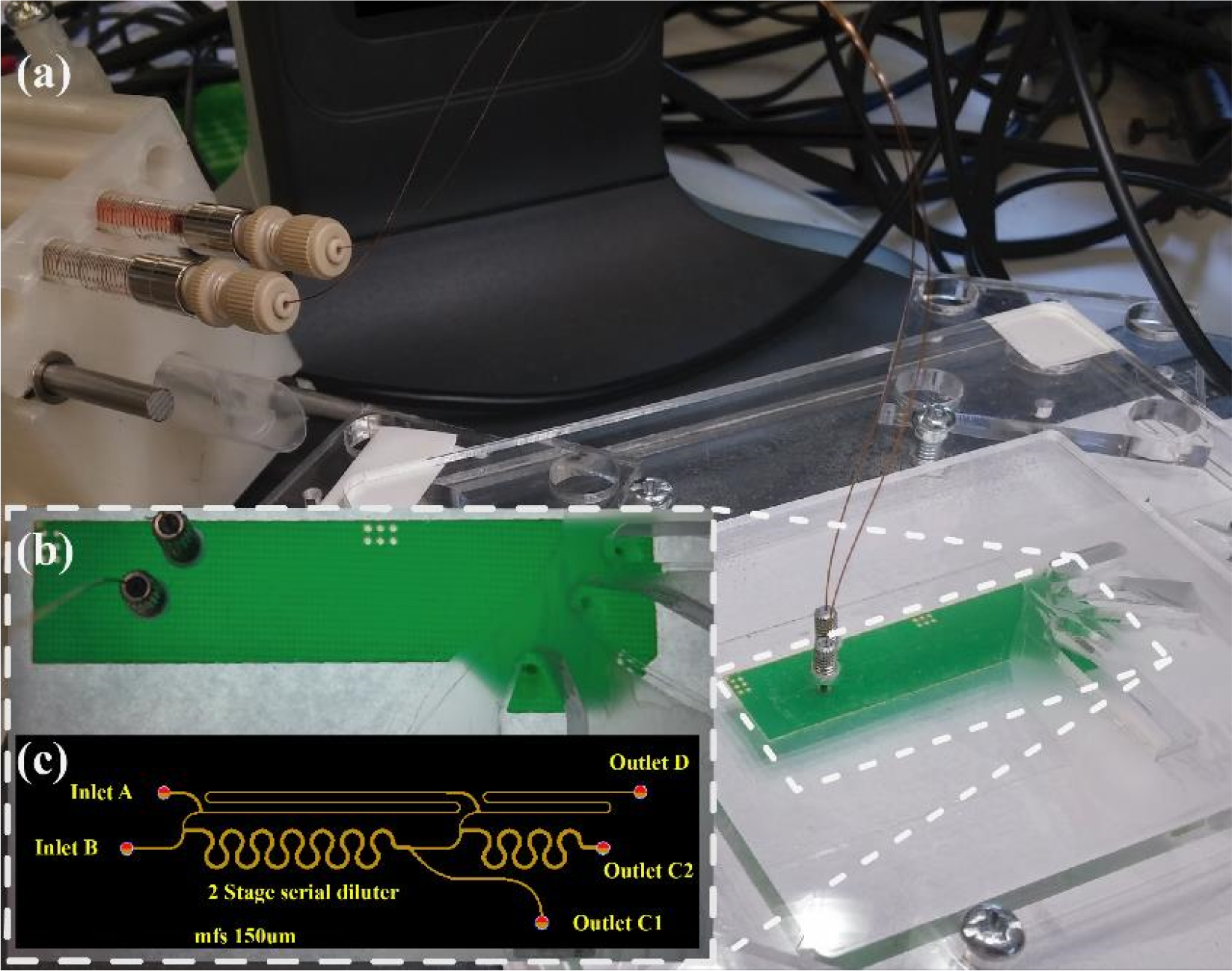
Experimental setup a) The prototyped 2-stage PCB diluter during performance validation experiment b) top view of the device c) design used for the prototype fabrication.

## 3. Results and discussion

### 3.1 Computational results

#### 3.1.1 First approximation

The first stage of the electrical network analogue of the microfluidic device illustrated in Figure 3b was used to define the required hydraulic resistances of the pressure and flow rate balanced modular unit cell. The network comprises 5 nodes (see Figure 3b positions 1 to 7) and 6 hydraulic resistances, R_1-2_ R_2-3_ R_2-6_ R_4-3_ R_3-5_ and R_5-7_. The desired DR of this unit cell has been selected to be equal to 2/3, while the nominal flowrate of the device was defined as 0.4μL/min after some iteration to facilitate sample formation at the outlets within a reasonable operating time (less than 3 minutes). Applying the mass conservation equation in nodes 2 and 3, we derive the following equations:The required dilution ratio (DR=2/3) complies with the following equation:

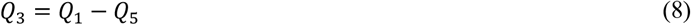

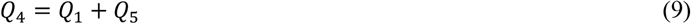

The flow rate balanced design satisfies the following 2 equations

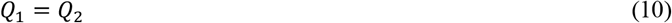

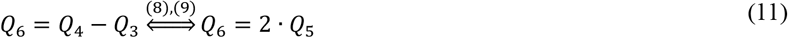

The required dilution ratio (DR=2/3) complies with the following equation

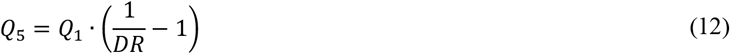

From equations (8) to (12) with the desired device specifications (DR and nominal flow rate), all flowrates in the modular unit cell can be computed. The minimum feature size of this technology is 150μm (MFS=150 μm) due to the DPR technical specifications. Both inlet microchannels were initially assumed to have the same channel width equal to double the MFS, i.e. *cw*_1-2_ = *cw*_4-3_ = 2 MFS = 300 μm. The channel width of the microchannel between positions 2 and 3 was selected to be slightly wider than the MFS, *cw*_2-3_ =160 μm while its length was selected arbitrary as L_2-3_=2.5mm. The mixing channel between position 3 to 5 was selected to be wider than the inlet channels since the flowrate though it is 50% higher than the nominal flowrate (DR=2/3), *cw*_3-5_ =380 μm. Equation 1 can be used as a rough estimate of the mixing zone channel length. The employed mixing length was increased by 30% as safety factor resulting in 58mm. The bypass diluent microchannel between positions 2 and 6 was defined with width *cw*_3-5_ =170 μm, while the outflow channel was selected initially to have a width of 300 μm as the device inlets. However, after the optimisation study the final width was defined to be 220μm, so that the resulting channel length facilitates the sampling position to be at certain distance from the dilution network (facilitating the device interfacing).

The pressure and flowrate balanced design approach together with the requirement for low pressure drop is defined by the following two equations.

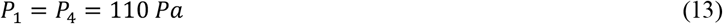

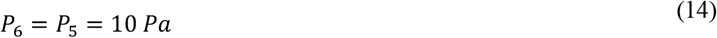

Using equation (3) the hydraulic resistances of branches *R*_*2-3*_ and *R*_*3-5*_ can be calculated. The four remaining hydraulic resistances (*R*_*1-2*_, *R*_*4-3*_, *R*_*2-5*_ and *R*_*5-7*_ can be determined after solving the following linear equations system. These equations are complying with the conservation of energy law along closed paths through the microfluidic networks (in accordance with Kirchhoff’s voltage law).

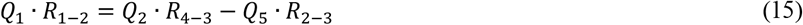

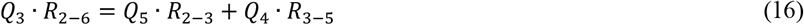

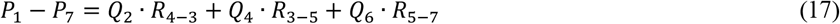

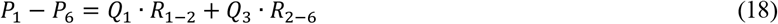

This methodology can be applied to design pressure and flowrate balanced modular unit cells of various dilution ratios by amending the DR parameter. The hydraulic network illustrated in Figure 2b can be designed in the same way if the hydraulic resistance R_5-7_ equals zero.

After obtaining the values of the hydraulic resistances, using equation (3) the microchannel lengths can be calculated. A Matlab custom-made script has been written for the solution of the system shown in eq (15) to (18) and the computation of the various hydraulic resistances and microchannel lengths (eq (1) to (3)). Matlab numerical results from the first cut approximation method are shown later together with COMSOL optimisation results.

#### 3.1.2 Simulation and optimisation

##### Single stage diluter

Figure 6 summarises the computational results of the device dilution performance when the flow rate through both inlets (A and B) equals 0.4 μL/min. This is the nominal flow rate of the device that offers sufficient time for the diluted sample to be formed at the device outlet (less than 3 minutes, since the optimised unit cell volume is 2.32 μL), i.e. steady state. The concentration field in Figure 6a verifies that the 6 double-loop micromixing channel has adequate length to achieve uniform mixing at the device outlet. Further refinement of the computational results is presented in Figure 6b and c. Figure 6b shows details around the merger point 3 where iso-concentration curves of diluted species with concentrations between 0.00 mol/L and 1.00 mol/L and step of 0.05 mol/L. Likewise, Figure 6c illustrates the iso-concentration curves of diluted species between 0.660 and 0.680 mol/L with a step of 0.002 mol/L along the 3^rd^, 4^th^ and 5^th^ double-loop microchannel, i.e. a finer range around the expected theoretical value (i.e. 0.667 mol/L). The iso-concentration lines in Figure 6b highlight that the two flow streams (sample and diluent) form a diffusion interfacial plane at approximately 2/3 of the microchannel width. After the merging point 3, the first 5 double loops of the mixing zone (as depicted in Figure 6c) result in a concentration field within a 0.002 mol/L values range between 0.666 and 0.668 mol/L.

**Figure 6:**
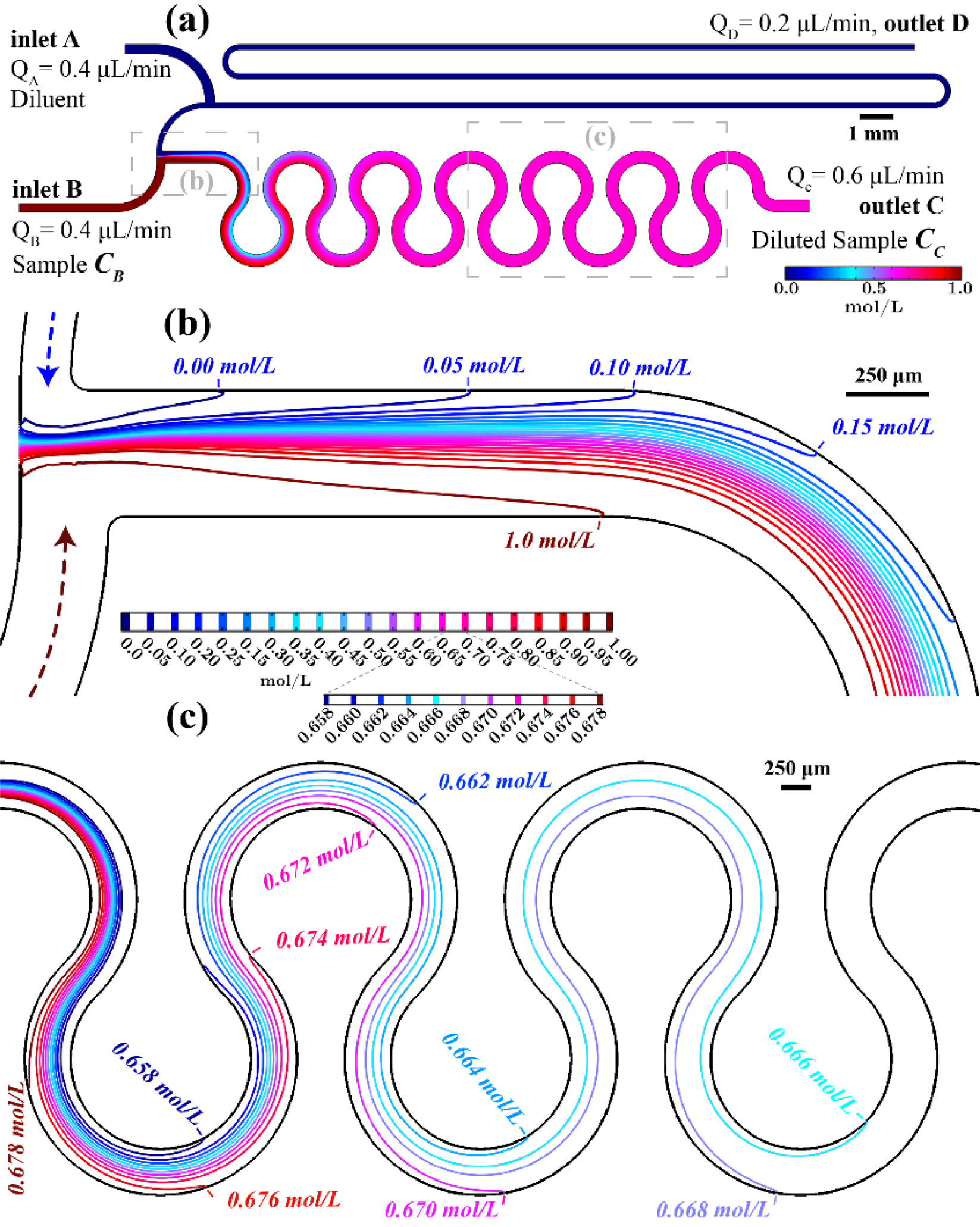
Pressure and flow rate balanced unit cell simulation results on the device symmetry x-y plane (z = 32 μm) **a)** concentration field overview **b)** iso-concentration lines at the merging point **c)** iso-concentration lines before the outlet of the mixing zone.

In Figure 7b the concentration field at the outflow surface for inlet flow rates of 0.2 μL/min, 0.4 μL/min, 0.6 μL/min and 0.8 μL/min is shown. The grid represents the spatial discretisation applied at the outflow surface. The ME is also reported for every instance proving that the optimised design generates uniformly mixed diluted sample at the nominal flow rate of 0.4 μL/min. For a lower flow rate (0.2 μL/min) the ME is identical (0.9984). In addition, the table presented in Figure 7a shows the pressure at the inlet surfaces for various flow rates, while the iso-pressure lines along the unit cell illustrate the positions experiencing equal pressure values. Evidently, the pressure balanced optimised design results in equal pressure formation on both inlets for every simulated flow rate. More specifically, the nominal inlet flow rate (0.4 μL/min) through both inlets generates 82.5 Pa or 8.4 mm H2O (mm water column). This order of pressure magnitude can be easily provided by a “burst blister” packaging. Furthermore, the pressure created on both inlet surfaces increases in proportion to the flow rate while remaining equal. Consequently, this design offers dilution rate performance that is tolerant of flow source instabilities. However, as indicated in Figure 7b for flow rates higher than 0.6 μL/min the ME drops (ME= 0.9838 for Q_A,B_= 0.8 μL/min).

**Figure 7:**
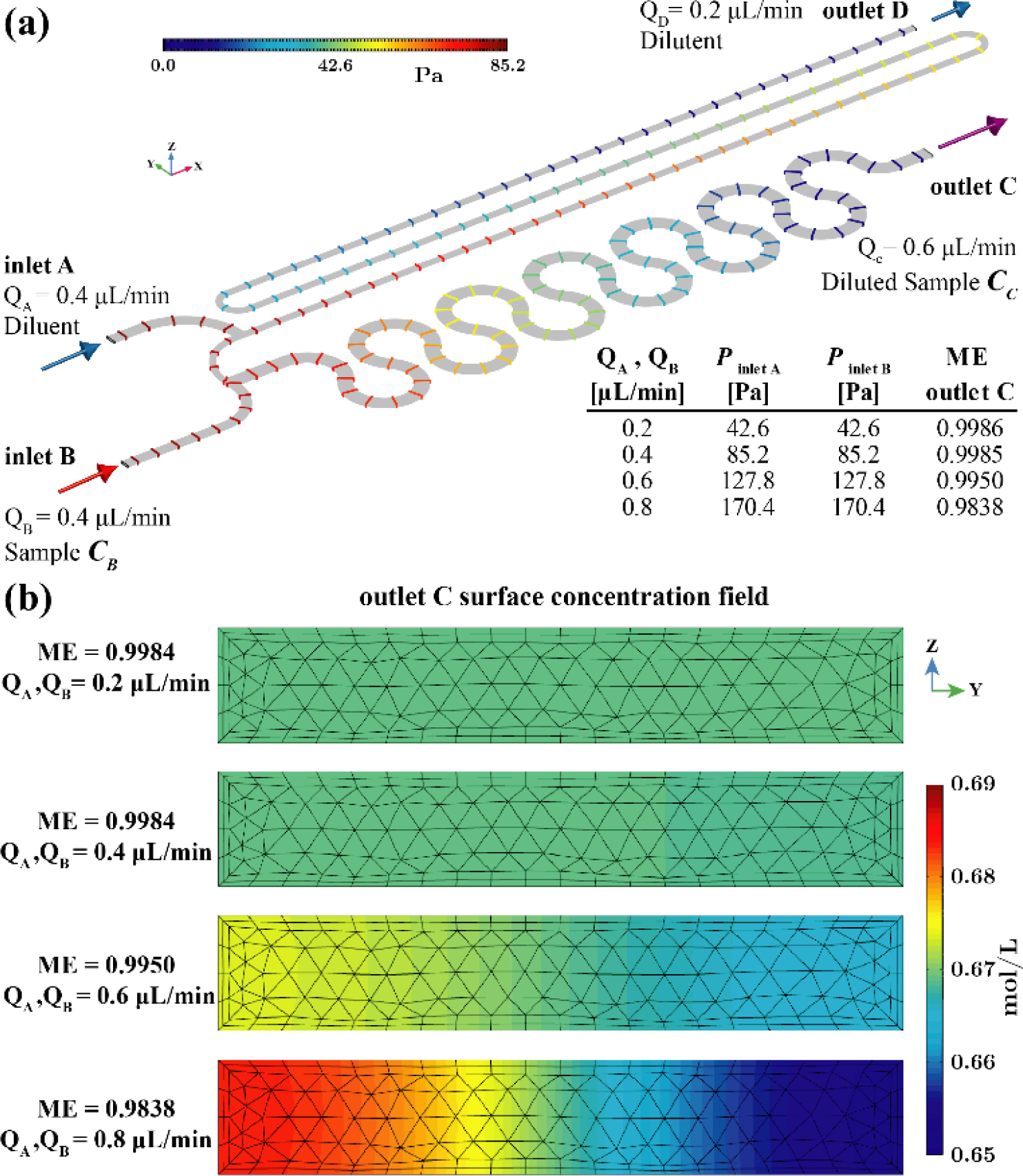
Pressure and flow rate balanced unit cell simulation results. a) iso-pressure lines b) concentration field on the outlet C surface for various flow rates

##### The 2-stage serial diluter

The optimised unit cell presented in Figure 2 can be considered as the fundamental design of a modular unit cell, the building block for multistage serial dilution networks. Figure 3 includes the resulting pressure and flow rate balanced 2-stage serial diluter after optimisation. The highlighted microfluidic network in Figure 3a is the modular unit cell that can be cascaded to form an n-stage serial dilution network as illustrated by the analogous electric circuit in Figure 3b.

In Figure 8a the reader can find computational results for the dilution performance when the nominal flow rate (0.4 μL/min) is applied through both inlets (A and B). The volume of the PCB-based 2-stage diluter is 3.87 μL and consequently less than 5 minutes are required to generate the two diluted sample concentrations (i.e. *C*_*1*_= 2/3 *C*_*B*_ and *C*_*2*_= 2/3 *C*_*1*_). In order to minimise the device footprint (105 x 8 mm^2^) the 2^nd^ stage mixing zone (between point 3’ and 5’) comprises a 3 double-loop microchannel. As depicted in Figure 8a such micromixing channel results in a uniformly mixed diluted sample at the *C*_*2*_ device outlet. Figure 8b and c show the iso-concentration curves of diluted species (glucose molecules) in water with concentrations between 0.00 mol/L and 1.00 mol/L with a step of 0.05 mol/L as they are formed after Stage 1 and Stage 2, merging points 3 and 3’ respectively. Similar to Figure 6b, the iso-concentration lines in Figure 8b and c show that the two flow streams (sample and diluent) form a diffusion interfacial plane at around the 2/3 of the microchannel width.

**Figure 8:**
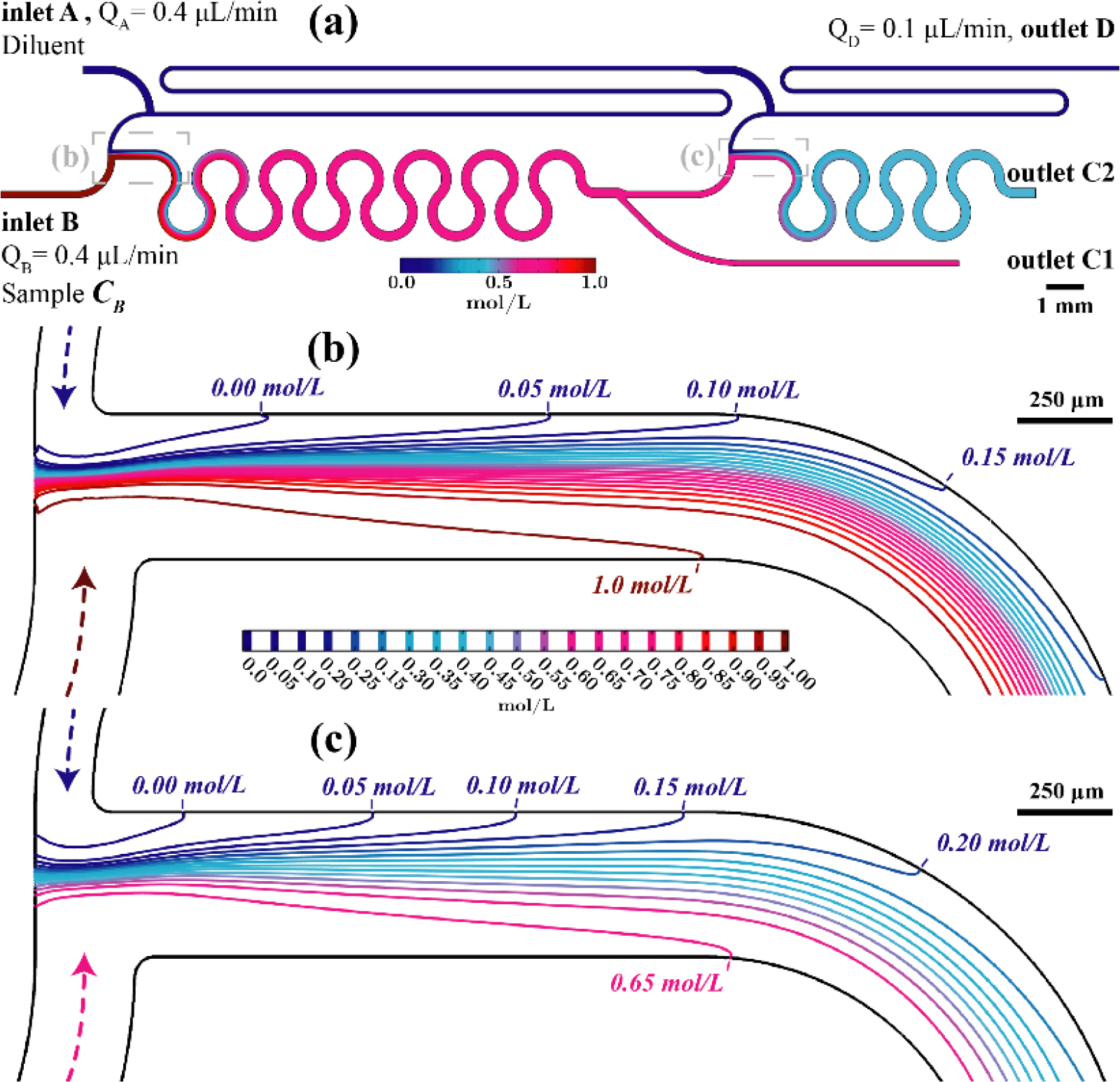
Pressure and flow rate balanced 2-stage serial diluter simulation results on the device symmetry x-y plane (z=32 μm) a) concentration field overview b) iso-concentration lines at the Stage 1 merging point c) iso-concentration lines at the Stage 2 merging point

Finally, Figure 9a shows the pressure drop along the 2-stage diluter, while Figure 9b i to iv illustrate the concentration field on the surface of outlets *C*_*1*_ and *C*_*2*_ for inlet flow rates 0.2 μL/min, 0.4 μL/min, 0.6 μL/min and 0.8 μL/min. Additionally, the inclusive table of Figure 9a summarises the computational results for the pressure on the inlet surfaces for the various flow rates. The iso-pressure lines in Figure 9a along the 2-stage diluter visualise the positions where the pressure level is the same. Evidently, the optimised 2-stage design results in equal pressure on both inlets for every flow rate. It was computed that pressure equal to 109.8 Pa is required to achieve the nominal inlet flow rate (0.4 μL/min) through both inlets. Due to the design optimisation objectives, if the applied pressure on both inlet surface increases, the flow rate increases proportionally while the generated diluted samples concentrations (*C*_*1*_ and *C*_*2*_) remain unaffected. However, for flow rates higher than 0.6 μL/min the ME drops, resulting in a non-uniform diluted sample concentration. Consequently, the dilution ratio is affected for both outlets. In detail, for the case of 0.8 μL/min inlets flow rate, the resulting *C*_*1*_ outlet sample concentration is *C*_*1*_= 0.68 mol/L. Thus, the generated dilution ratio is 0.68 instead of 0.67, which is the design dilution ratio of both stages (*C*_*1*_= 2/3 *C*_*B*_ or 0.67 *C*_*B*_). This minor deviation is attributed to the fact that the sampling microchannel (see Figure 3a, between pos. 5 to 7) at position 5 is placed on the same side as the sample inlet (inlet B). Consequently, the non-uniformly generated diluted sample gives a higher concentration on the inlet B side. The 2^nd^ Stage is provided with lower concentration inlet sample and the diluted sample through outlet *C*_*2*_ presents lower concentration (i.e. *C*_*2*_= 0.43 mol/L instead of 0.44 mol/L in case of nominal inlet flow rate).

**Figure 9:**
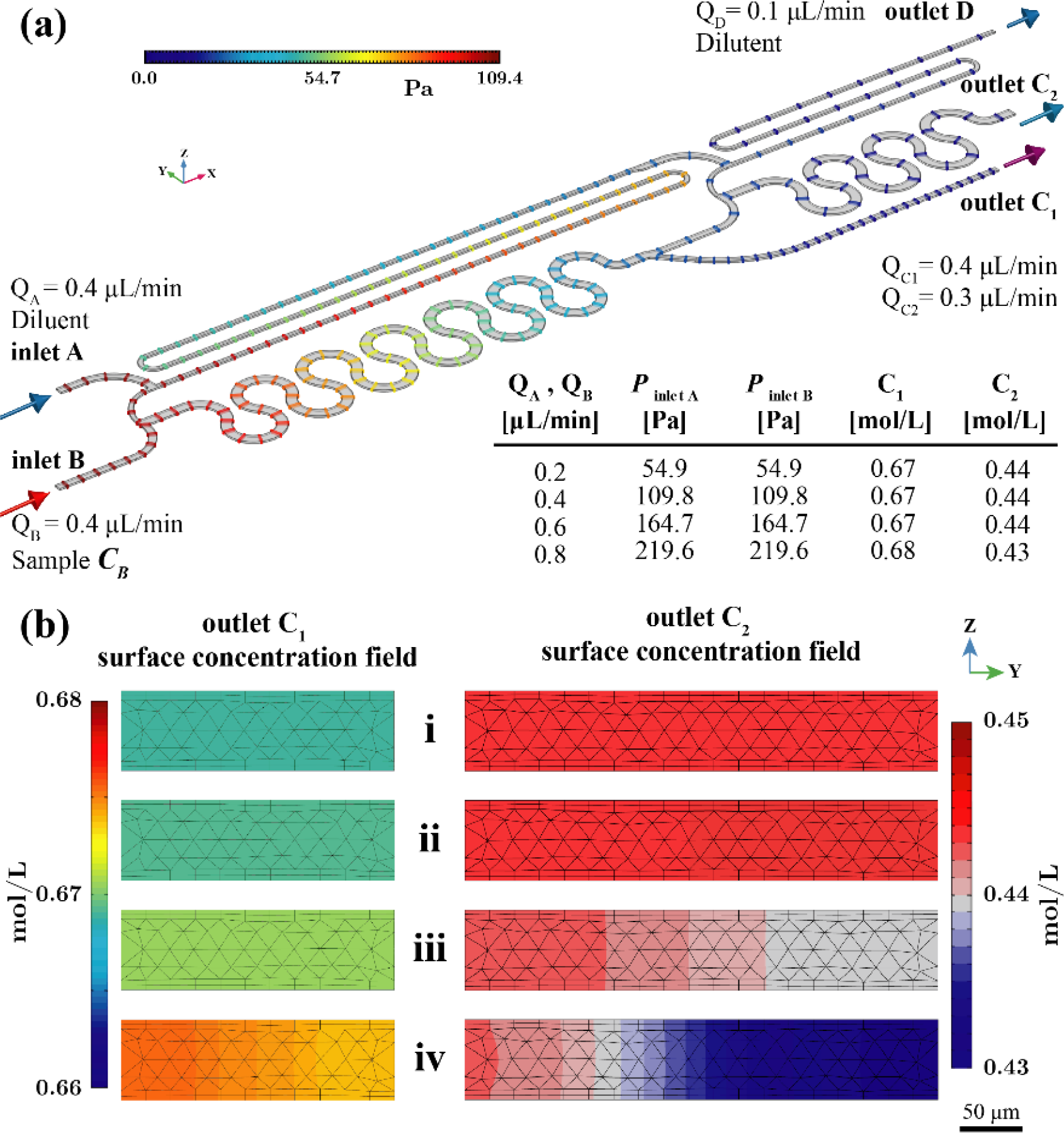
Pressure and flow rate balanced 2-Stage serial diluter simulation results. a) iso-pressure surfaces b) concentration field on the outlets *C1* and *C2* surface for various flow rates.

In Table 1, columns 2 and 3, we summarise the results of the “first-cut” approximation numerical simulation based on the Matlab code. These microchannel lengths are not the final, fabricated ones. These values were used to define an initial design of the network that was then used in an automated optimisation study using Solidworks and COMSOL Multiphysics in a coupled manner. The final geometrical parameters of the modular unit cell (Table 1, columns 4 and 5) were used for the fabricated and characterised devices that will be presented later. To compare the analytical calculations based on equations (1) to (3) and (8) to (18) with the “first-cut” approximation numerical simulation, we used the optimised microchannel widths (Table 1, column 4) and the resulting pressure values of the simulation results as input to solve the system of equations. Table 1, column 6 presents the obtained microchannel lengths from Matlab. Comparing the microchannel lengths in column 5 and 6 in Table 1, it can be seen that the first-cut approximation method in Matlab is in good agreement with COMSOL.

**Table 1.** Modular pressure and flow rated balanced unit cell. Microfluidic network geometrical parameters results.

| Channel | Matlab First cut approximation<br>P1-P6 = 100 Pa, P1 = 110 Pa |  | Comsol optimisation results<br>P1-P6 = 85.4 Pa, P1 = 109.8 Pa |  | Matlab Validation results<br>P1-P6 = 85.4 Pa, P1 = 109.8 Pa |
| --- | --- | --- | --- | --- | --- |
| | Width $cw$<br>[$\mu\text{m}$] | Length $L$<br>[mm] | Width $cw$<br>[$\mu\text{m}$] | Length $L$<br>optimization<br>study<br>[mm] | Length $L$<br>validation calculations<br>[mm] |
| 1-2 | 2 MFS = 300 $\mu\text{m}$ | 16.0 mm | 2 MFS = 300 $\mu\text{m}$ | 3.7 mm | 3.4 mm |
| 4-3 | 2 MFS = 300 $\mu\text{m}$ | 18.7 mm | 263 $\mu\text{m}$ | 5.4 mm | 5.3 mm |
| 2-3 | 160 $\mu\text{m}$ | 2.5 mm | 160 $\mu\text{m}$ | 2.5 mm | 2.5 mm |
| 2-6 | 170 $\mu\text{m}$ | 69.0 mm | 170 $\mu\text{m}$ | 67.5 mm | 69.0 mm |
| 3-5 | 380 $\mu\text{m}$ | 57.9 mm | 380 $\mu\text{m}$ | 57.9 mm | 57.9 mm |
| 5-7 | 2 MFS = 300 $\mu\text{m}$ | 8.5 mm | 220 $\mu\text{m}$ | 14.4 mm | 14.5 mm |

Finally, Figure 10 summarises the resulting design rules of the modular pressure and flow rate balanced unit cell of Figure 2b for various dilution ratios, ranging from 2:3 to 10:11, based on the custom-made Matlab script. All the plotted solutions were derived with constant channels widths, equal to the computationally optimised ones (see Table 1 column 4).

**Figure 10:**
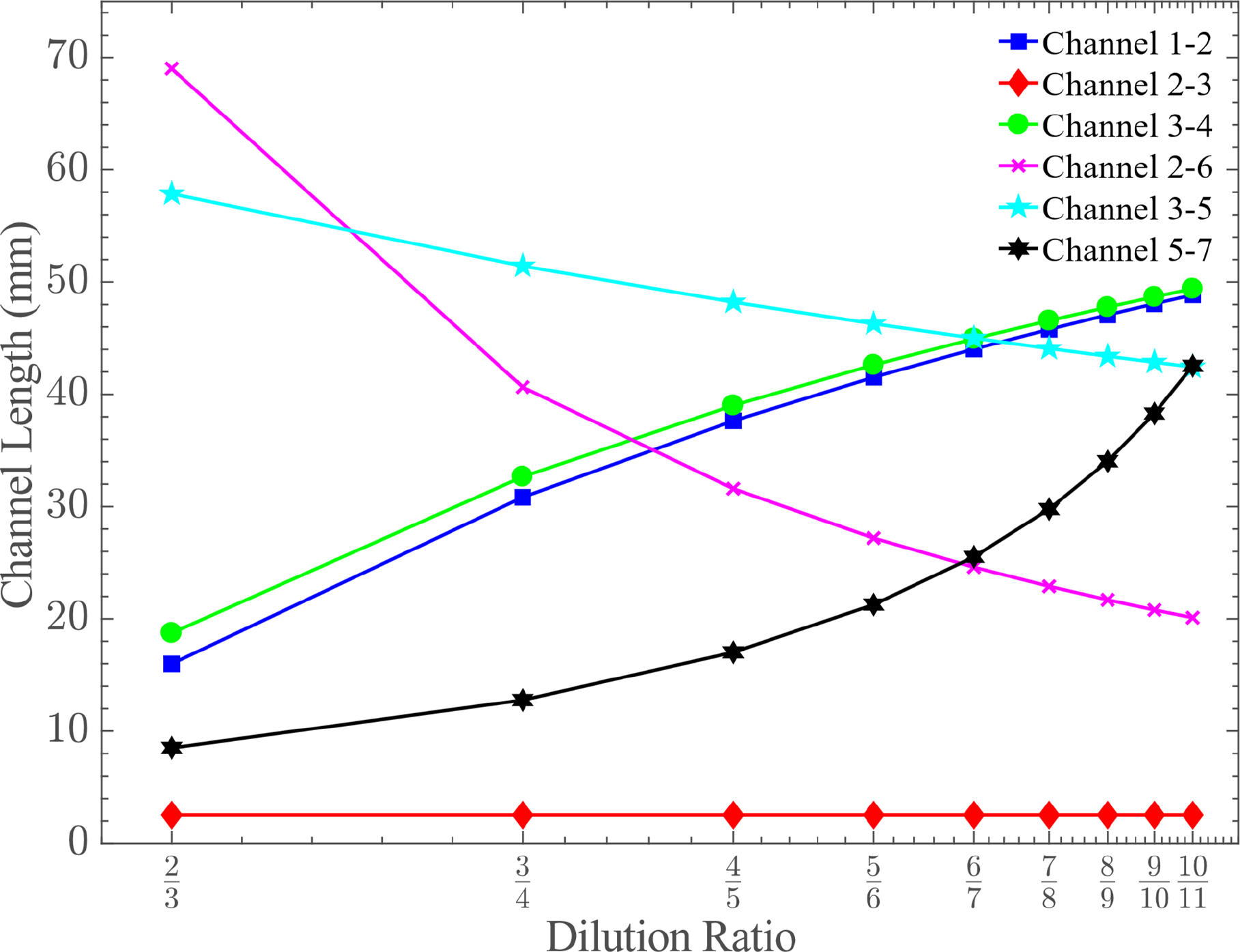
The design rules of the pressure and flow rate balanced modular unit cell showing the required microchannel lengths for various dilution ratios. Microchannel widths are kept the same as the optimised modular unit cell generating dilution ratio 2:3.

### 3.2 Experimental performance and validation

The device dilution ratio performance was validated using a commercial glucose meter (Accu-Check Aviva). In detail, 0.1 M Phosphate-buffered saline (PBS) (inlet A) and 15 mM glucose (Sigma-Aldrich) in PBS (inlet B) were prepared as diluent and sample solutions, respectively. A syringe pump with two 200 μL syringes supplied the 2-stage PCB serial diluter inlets with sample and diluent solution maintaining the flow rate through both inlets equal and constant. Two different flow rates were applied, i.e 0.2 μL/min and 0.4 μL/min. Each of the generated diluted samples (*C*_*1*_ and *C*_*2*_) was measured three times. In Figure 11 the statistically analysed dilution ratio measurements are presented (rectangular and triangle marks for the two applied flow rates), while the design relevant triangular mark pointing downwards is provided for comparison.

A highlighted area marking the acceptable measurement error due to the glucose meter accuracy (±10%) is also provided. The average measured dilution ratio is presented for the different device outlets. The error bars indicate the standard deviations for each measured sample. In the case of the 0.2 μL/min inlet flow rate, the 1^st^ and 2^nd^ stage dilution ratio was found to be equal to 0.65 ± 0.02 and 0.46 ± 0.01, in good agreement with the design values, taking also into consideration the glucose meter accuracy. Likewise, for the nominal flow rate (0.4 μL/min) the dilution ratio performance of both stages was found equal to 0.70 ± 0.01 and 0.43 ± 0.01, respectively.

The sampling point 5 (see Figure 3a) is on the same side as inlet B (point 4). If the flowrate through the micromixing zone (between point 3 and 5) is higher than assumed in the design calculations (*Q*_*4*_ = *Q*_*1*_/*DR* = 0.6 μL/min) the microchannel length does not allow for uniform mixing of sample and diluent. In parallel, inlet B is used as sample inlet and consequently, the sample stream flowing along the sidewall of the micromixing zone (that is on the same side with the sampling channel, branch 5 to 7 in Figure 3) is not fully diluted. Therefore, the liquid flowing through the outlet C1 is expected to have higher concentration than designed. The table in Figure 9a shows that for the case of Q_A_, Q_B_=0.8μL/min, the resulting concentration in outlet C_1_ is 0.68mol/L instead of the 0.67mol/L. Additional experiments with higher flow rates (i.e. 0.8 μL/min) resulted in dilution performance outside the design values, as predicted by the simulation results shown in Figure 9.

**Figure 11:**
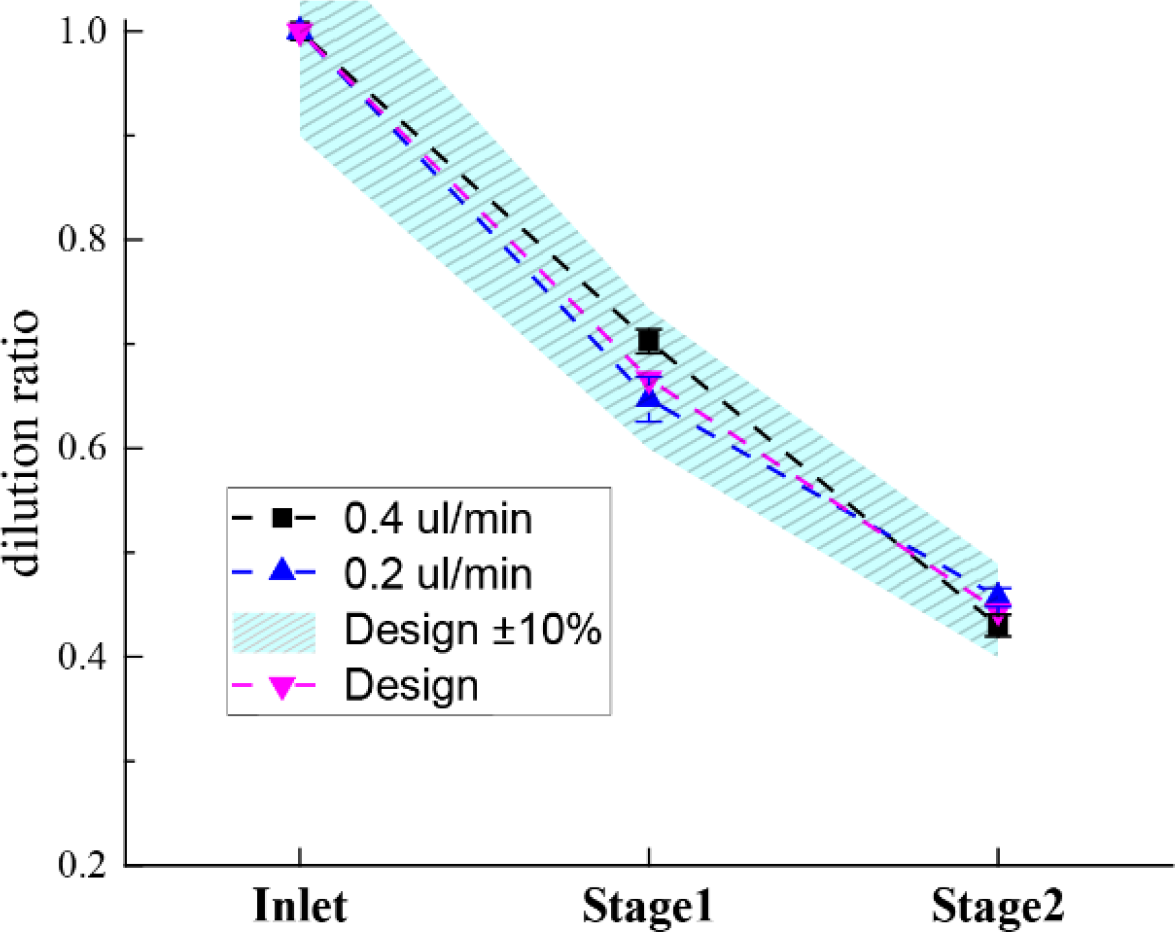
Dilution ratio validation of the 2-stage PCB prototype using commercial Accu-Check^®^ glucose meter. The solid marks represent the dilution performance under 0.4 and 0.2 μL/min flow rate concentration measurements.

## 4. Conclusions

In this work a modular serial dilution unit cell was conceived, designed, optimised and fabricated, suitable for LoPCB platforms, offering both, pressure and flow rate balance. The presented, indicative, unit cell performs 2:3 sample dilutions and was used to form a 2-stage step wise serial dilution network generating two diluted samples (i.e. 2:3 and 4:9 diluted inlet sample). The prototyped design comprising the modular unit cell was optimised using COMSOL Multiphysics^®^ with a laminar flow and diffusion-advection model. The final 2-stage PCB microfluidic device was fabricated using standard, commercially available PCB manufacturing processes and the performance was validated experimentally using glucose. Simulation results were in good agreement with measured results.

If required, the modular serial dilution unit cell can be repeated in a cascade manner to form multistage serial dilution networks of compact footprint (around 75 x 8 mm^2^ per stage), while the pressure and flow rate balanced capability minimises the number of flow sources. Both diluent and sample can be driven through the device using a single flow source. In detail, it can operate with a single pressure source (compressed air chamber or blister packaging) or a constant flow rate source (syringe pump). This allows the device to be used as part of a simple PoC test requiring serial dilutions for calibration. However, the trade-off for the modular pressure and flowrate balanced design is that the flowrate of diluted sample and diluent entering each subsequent stage is much lower than the previous, i.e. 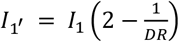 resulting in extended time for the diluted samples appear at the outlets.

The low pressure required on both inlets (109.8 Pa or 11.2 mmH_2_O) ensures the above. In addition, the generated dilution ratios tolerate flow source instabilities, since the dilution ratio remains constant as long as the nominal flow rate or inlet pressure is not exceeded.

Taking all the above into consideration, the proposed design could be an ideal candidate for affordable PoC tests relying on quantitative assays, where, for example, serial dilutions of a known sample concentration are required so that a standard curve can be used as a reference. The modular unit cell can be cascaded for more complicated multistage serial dilution network applications and is highly compatible with PCB-based biosensors and electronic components, allowing for the monolithic integration of all of them on PCB, once a complete LoPCB platform for quantitative PoC medical diagnostic tests is sought after.

## Acknowledgements

The authors wish to acknowledge the financial support of the A.G. Leventis Foundation and EPSRC EP/L020920/1. We also thank Newbury Electronics Ltd. (Faraday Road, Newbury, West Berkshire RG14 2AD, UK) for their valuable input in manufacturing the presented PCB microfluidic devices and Dr Despina Moschou for helpful discussions.

